# Identity-by-descent (IBD) segment outlier detection in endogamous populations using pedigree cohorts

**DOI:** 10.1101/2024.08.07.607051

**Authors:** Shi Jie Samuel Tan, Huyen Trang Dang, Sarah Keim, Maja Bućan, Sara Mathieson

## Abstract

Genomic segments that are inherited from a common ancestor are referred to as identical-by-descent (IBD). Because these segments are inherited, they not only allow us to study diseases, population characteristics, and the sharing of rare variants, but also understand hidden familial relationships within populations. Over the past two decades, various IBD finding algorithms have been developed using hidden Markov models (HMMs), hashing and extension, and Burrows-Wheeler Transform (BWT) approaches. In this study, we investigate the utility of pedigree information in IBD outlier detection methods for endogamous populations. With the increasing prevalence of computationally efficient sequencing technology and proper documentation of pedigree structures, we expect complete pedigree information to become readily available for more populations. While IBD segments have been used to *reconstruct* pedigrees, because we now have access to the pedigree, it is a natural question to ask if including pedigree information would substantially improve IBD segment finding for the purpose of studying inheritance. We propose an IBD pruning algorithm for reducing the number of false positives in IBD segments detected by existing software. While existing software already identify IBD segments with high success rates, our algorithm analyzes the familial relationships between cohorts of individuals who are initially hypothesized to share IBD segments to remove outliers. Our algorithm is inspired by a k-Nearest Neighbors (kNN) approach with a novel distance metric for pedigrees with loops. We apply our method to simulated genomic data under an Amish pedigree, but it could be applied to pedigrees from other human populations as well as domesticated animals such as dogs and cattle.

## Introduction

Studying pedigree structures has proven to be very useful for estimating mutation and recombination rates in humans and other species [7]. Pedigrees not only contain significant information for the study of short-term evolution in natural populations [9] but also help us to better understand disease heritability [3].

The pedigrees of endogamous populations are particularly helpful, as endogamy is characterized by marriage within a small group for geographic, religious, or cultural reasons. These pedigrees and associated genomic data allow us to attribute genetic variants and related traits to specific parts of the genome with greater confidence [22, 27]. Even when the pedigree size is small and genetic information is relatively minimal, studying genetic inheritance within endogamous populations can help researchers better understand common diseases [15, 32].

IBD segments refer to portions of genomic sequence inherited from a recent common ancestor within a population [33]. As these segments are passed down through generations, they provide insights into the characteristics of a population and the distribution of rare and moderately common genetic variants. Additionally, they can aid in uncovering obscure familial connections within populations [29, 33]. In spite of the numerous applications that IBD segments provide, finding IBD segments is known to be a computationally challenging task. Because IBD finding often has to compare pairs of individuals, naive IBD detection algorithms tend to scale quadratically with the number of individuals. Hence, pattern matching and other matching techniques were often not computationally efficient [31]. In 2007, Purcell *et al*. demonstrated that it is possible to infer relatedness between individuals using hidden Markov model (HMM) techniques on a dense set of markers [2, 28]. By using HMMs, Purcell *et al*. developed `PLINK` to estimate the underlying IBD state between pairs of individuals. Other algorithms like `Beagle` [5] have since been developed with similar HMM techniques to use a probabilistic approach to identify IBD segments.

While HMM-based algorithms can detect IBD segments with high resolution, they have also been shown to be computationally inefficient for large cohorts since they have to check all pairwise relationships [16]. In 2008, Gusev *et al*. proposed `GERMLINE` [17], a new type of IBD segment finding algorithm that scaled to very large populations. It takes advantage of the efficiency of hashing functions and creates hash tables between short haplotype matches before extending them into IBD segments [31]. The technique of hashing and then extending has been utilized and improved by various IBD detection algorithms like `iLASH` and `FastSMC` [29, 30]. In fact, `FastSMC` first leverages `GERMLINE`’s hashing and extension algorithm to identify IBD segment candidates before using an HMM for validation. This suggests the possibility of combining different IBD detection techniques to maximize the accuracy of IBD finding.

Other than using HMM or the hashing and extension technique, another commonly applied type of IBD detection algorithm uses variations of Burrows-Wheeler Transform (BWT). Even though BWT was used for aligning genetic sequences [21], Durbin proposed a novel approach using positional Burrows-Wheeler Transform (PBWT) to find all maximal haplotype matches for IBD detection in 2014 [12]. Durbin’s approach first converts the data set into a different representation based on positional prefix arrays before finding matches. PBWT then inspired other variations of PBWT-based approaches like `phasedibd` and `Hap-IBD` which use templated positional Burrows-Wheeler Transform (TPBWT) and error-adjusted PBWT respectively [31].

In our previous study, we presented a novel algorithm, `thread`, for reconstructing ancestral haplotypes given an arbitrary pedigree structure and genotyped or sequenced individuals [13]. Our algorithm uses an IBD detection software as a subroutine to identify IBD segments belonging to a population before utilizing the identified IBD segments to reconstruct ancestral haplotypes. We applied `thread` to an extended Old Order Amish pedigree [26] with 1338 individuals [1, 14]. Our algorithm used the genetic data from 394 genotyped individuals to reconstruct the ancestral haplotypes of the larger pedigree to study inheritance patterns of mood disorders. The way we designed `thread` allows it to be effective even when there are loops, inter-generational marriages, and remarriages within the pedigree. Pedigree loops occur through the marriage of close relatives. The results of our study showed that the reconstruction success rate for `thread` is related to the ability of the IBD detection software to correctly identify an IBD segment and the group of individuals that shares the segment by inheritance.

In this study, we explore the ways that we can utilize information from the pedigree structure to improve the performance of IBD detection software for endogamous populations. Even though current IBD finding algorithms are able to correctly identify IBD segments with high success rates, they may not have accurately determined the group of individuals that actually inherited the IBD segment recently. In other words, there may be outliers that possess the IBD segment by chance and did not actually inherit the IBD segment, especially in endogamous populations. These segments would be identical-by-state (IBS), which is sometimes referred to as background IBD. A natural question is whether we can utilize pedigree information to improve our ability to identify IBD segments and correctly determine the groups of individuals that share them. A more accurate set of IBD calls would advance our haplotype reconstruction algorithms such as `thread`. Researchers would also be able to better determine the extent of regions that harbor risk alleles for both common and rare diseases. By using the information from the Amish pedigree examined in our previous work, we identify ways in which understanding relationships between individuals that share IBD segments could potentially improve the accuracy of IBD calls. Because endogamous populations such as the Amish have complex relationship structures like loops and inter-generational marriage, studying their pedigree allows us to better understand how knowledge of kinship affects IBD detection.

Here we present a novel algorithm that uses the pedigree structure to remove individuals who were wrongly inferred to share IBD segments by an existing IBD detection algorithm. Our method is inspired by the k-nearest neighbors (kNN) algorithm [18] and uses the shortest path between individuals in the pedigree as a distance metric to determine outlier status. Compared to `HaploScore` [11] that was introduced by Durand *et al*. in 2014, our algorithm focuses on improving `phasedibd` instead of `GERMLINE` and does not require estimates of the genotyping error rate and the switch error rate. However, the success of our algorithm hinges on judicious choices for threshold values that we define in the following sections. Our results show that our algorithm successfully improves the ratio of true positive individuals to false positive individuals for groups of individuals that were identified to have shared the same IBD segment by existing IBD detection software. Our algorithm can be applied to use cases that require conservative estimates of individuals that have inherited particular genetic segments. This work represents a key step toward understanding the use of pedigree information in IBD detection, and could be applied beyond humans to other species with pedigree information. Our code is available open-source: https://github.com/mathiesonlab/ped-aware-knn.

## Materials and Methods

### Problem statement

The first input to our proposed algorithm is a pedigree structure *P*. Similar to our previous work in [13], we denote each individual *p* ∈ *P* (aside from founders and married-in individuals). For each *p* ∈ *P*, we have information about the mother and father denoted as *p*^(*m*)^ and *p*^(*f*)^ respectively. For the founders and married-in individuals in *P*, we denote their parents *p*^(*m*)^ and *p*^(*f*)^ as 0’s. The pedigree can often be thought of as a directed graph that may contain loops. As mentioned formally in [4, 8], we observe a loop in the pedigree when the undirected version of the directed marriage graph contains a cycle. Loops usually exist when the *p*^(*m*)^ and *p*^(*f*)^ of a particular *p* ∈ *P* share a common ancestor *p*^*′*^ that is also in *P*.

The second input is a set of IBD segments that correspond to genetic information shared by pairs of individuals that belong to the pedigree. This set of IBD segments comes from a user-determined IBD finding software like `GERMLINE` or `phasedibd` after providing the software with haplotype data. Note that we refer to pairs of individuals, but the specific haplotype information is included as well.

The output of our kNN algorithm is a set of IBD segments that corresponds to genetic information shared by pairs of individuals that belong to the pedigree *P*. Note that the final number of pairs of individuals that share a particular IBD segment is less than or equal to the original number of pairs since our algorithm only *removes* individuals from the cohort. The goal is to use relationships between individuals to prune some of the IBD segments that are actually IBS or background IBD by removing the individuals that are falsely associated with a group of individuals that truly share the IBD segment.

### High level description

Our proposed kNN algorithm (shown in Figure 1) has two main steps: cohortization and filtration. In the cohortization step, we first consider each unique IBD segment independently. Note that our second input above only gives us a set of IBD segments that corresponds to *pairs* of individuals. Hence, we first identify the unique IBD segments before we group all the individuals that share the same IBD segment into cohorts. Subsequently, we proceed to the filtration step where we use our modified kNN algorithm to remove outliers from the cohorts.

**Figure 1.**
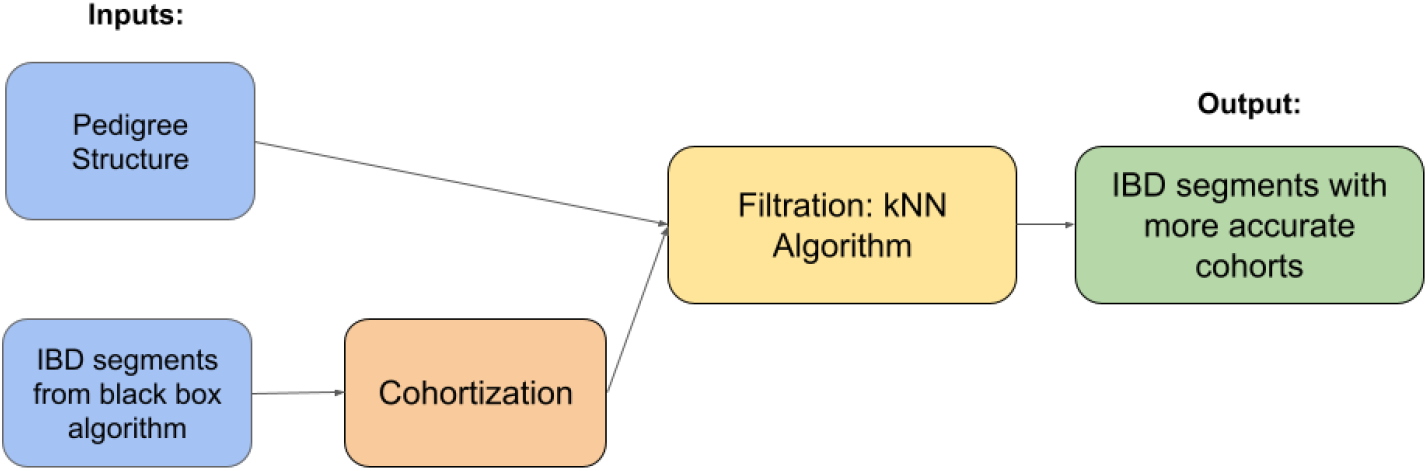
Workflow of our proposed algorithm. Pairwise IBD segments are obtained from an existing algorithm such as `GERMLINE` or `phasedibd`. We then group individuals into cohorts that share the same IBD segment. After filtering putative false positives, we are left with a potentially more accurate subset of IBD cohorts.

### Cohortization

After obtaining a set of IBD segments that correspond to pairs of individuals from the pedigree using an existing IBD detection algorithm, we adopt a cohort-based approach to remove false positives from the set of IBD segments. A cohort, in the context of our paper, is defined to be a group of individuals who are determined by the IBD detection algorithm to have shared a particular IBD segment [13].

However, the original cohorts do not necessarily represent the real set of individuals that share the same IBD segment. Individuals could be placed in a cohort due to sharing a segment through IBS or background IBD, especially if they are not closely related to the rest of the cohort. By removing the outliers based on the pedigree structure, our proposed algorithm is able to return cohorts that share IBD segments with higher accuracy.

### Toy example

In Figure 2, we provide a hypothetical situation to illustrate how to filter individuals who do not belong to the cohort. In this example, the software inferred an IBD segment that is shared across individuals 1, 2, and 4. However, since 1 and 2 are more closely related, it is very likely that the IBD observed in individual 4 is a false positive. Our kNN algorithm should detect that 4 is an outlier because individuals 1 and 2 are siblings (2 generations apart) whereas individual 4 is 4 generations away from both individuals 1 and 2.

**Figure 2.**
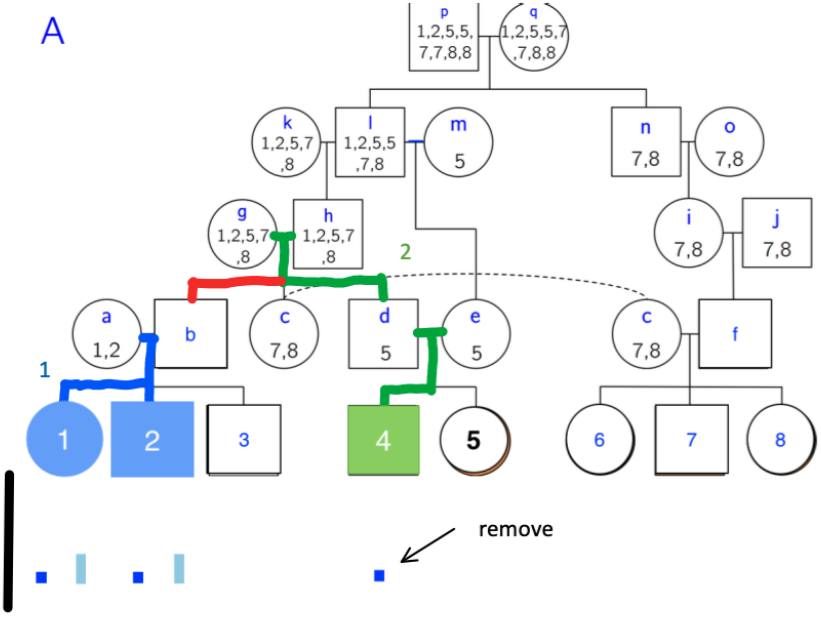
The black line represents an entire chromosome, and the blue segments represent IBD segments. Dark blue segments are inferred by IBD software, and light blue segments represents the ground truth. Our KNN algorithm removes the dark blue segment that belongs to individual 4.

### k-Nearest Neighbors Algorithm

#### Description of Algorithm

Our proposed k-Nearest Neighbors (kNN) algorithm is adapted from an outlier detection algorithm by Hautamaki [18]. One of our main contributions is the distance metric between individuals. Due to complex pedigree features such as loops, there are often multiple paths between pairs of individuals in this pedigree. In our algorithm, we utilized the shortest path between individuals (i.e. to the most recent common ancestor) as a distance metric *δ* for the k-Nearest Neighbors algorithm to identify the outliers in each cohort. Before we provide the pseudocode for our algorithm, we first provide some definitions and notations.

Let *k* ∈ ℕ be a user-defined parameter for the number of neighbors in the kNN algorithm. Let *C*_*i*_ be the *i*^th^ individual in the cohort *C*. Then, define *K*_*i*_ to be the set of *k* individuals in *C* that is the closest to *C*_*i*_. In other words, *K*_*i*_ should contain the *k* individuals in *C* whose shortest paths to *C*_*i*_ have the fewest edges compared to all other individuals in *C*. Define *L*_*i*_ to be kNN density of *C*_*i*_ such that

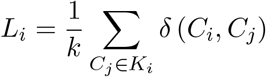

where *δ* (*C*_*i*_, *C*_*j*_) denotes the number of edges in the shortest path between individuals *C*_*i*_ and *C*_*j*_. Note that the set of paths that we consider when evaluating *δ* (*C*_*i*_, *C*_*j*_) only contains valid hereditary paths. In other words, we do not consider paths that connect two individuals via mutual descendants.

The algorithm computes *L*_*i*_ for all *C*_*i*_ in the cohort *C* and stores them in a list *L* in ascending order. In other words, *L* = {*L*_1_, *L*_2_, …, *L*_|*C*|_} where *L*_*i*_ ≥ *L*_*i*−1_. We re-index the individuals in the cohort so that *L*_*i*_ ∈ *L* now corresponds to the new *C*_*i*_. We now define *T* as the following:

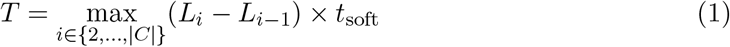

where *t*_soft_ ∈ (0, 1) is a user-defined parameter and set the soft threshold as

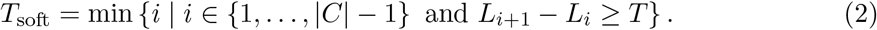

We can interpret *T*_soft_ as the index of the individual before we see a jump in kNN density. We consider that as an indicator that the individuals corresponding to indices that came after that index are potential outliers. To avoid the possibility of removing far too many individuals leaving us with far too few individuals in our cohort, we introduce the hard threshold *T*_hard_ which is defined as the following:

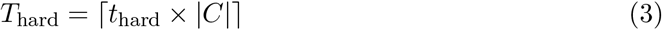

where the parameter *t*_hard_ ∈ [0, 1) is the minimum fraction of the cohort to keep.

Lastly, we let the final threshold be

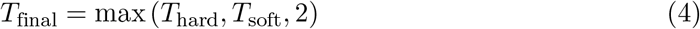

where we take the maximum of the two thresholds and make sure that the final cohort has at least two individuals. The algorithm outputs 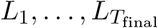 as the predicted true cohort and removes individuals 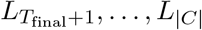. In other words, our algorithm now removes at most 1 − *t*_hard_ fraction of the original cohort. The values for the parameters, *k, t*_soft_ and *t*_hard_ can be chosen based on several different characteristics of the pedigree that include the cohort size, the number of cycles in the pedigree structure, etc. For example, a larger *k* value might be useful for larger cohort sizes. Likewise, we might consider using larger values for *t*_soft_ and *t*_hard_ if we want to keep most of the original cohort, i.e., a more conservative removal approach.

##### Algorithm 1

Pseudocode for our modified kNN algorithm.

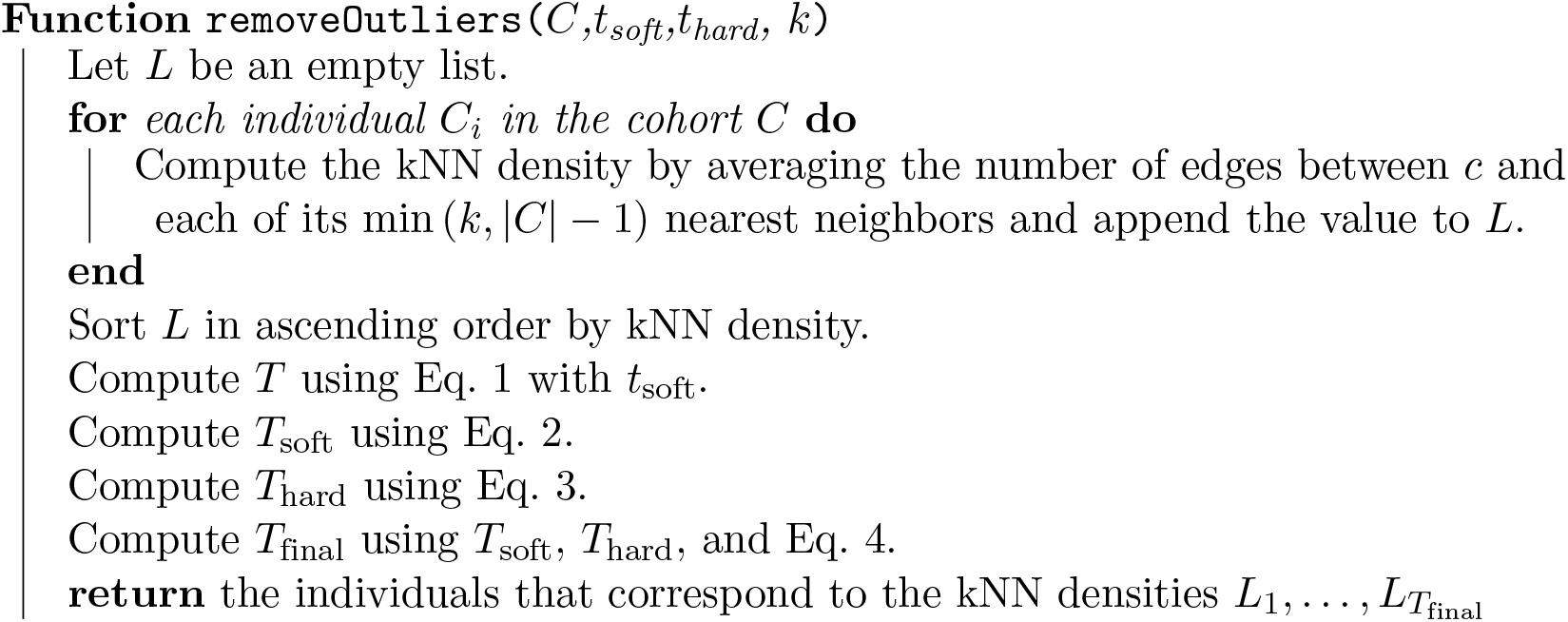

### Validation

Our goal for validation is to evaluate the effectiveness of our kNN algorithm at removing false positives. Because the removal of false positives often takes place at the expense of true positives, we also examine the cost of false positive removals as a secondary metric of success by studying the number of individuals that were falsely removed from a cohort when they, in fact, share an IBD segment with individuals from the cohort. To accurately assess the kNN algorithm, we compared the number of false positives in the cohorts constructed by `phasedibd` before and after the application of our kNN algorithm. We also performed a similar comparison between the number of true positives before and after applying our kNN algorithm to understand the cost of removing false positives.

Similar to the dataset we prepared in our previous work [13], we simulate a realistic endogamous population by first generating a founder population using `msprime` [20], with recombination rates drawn from the HapMap recombination map [10]. Then we simulated 25 generations of pre-migration endogamy using `Ped-sim` [6], to create founders for the North American Amish population. Finally, we again utilized `Ped-sim` to simulate the 1338 individuals under our known Amish pedigree structure, from which we retained genotype information for 394 descendants. We varied the genotyping error rate for our `Ped-sim` simulations over the following values 0.0, 0.005, and 0.01 [6]. This allows us to construct test cases for cohorts that share the same IBD segment using `Ped-sim`’s output to better evaluate our algorithm’s performance with respect to different genotyping error rates. Note that we assumed that `Ped-sim`’s IBD segment output is the ground truth. Then, we extracted the descendants that are originally genotyped and ran `phasedibd` on these descendants. In the Results section, we briefly compare `phasedibd` with `GERMLINE` and justify why we choose to use `phasedibd` for our study.

For each inferred or ground truth IBD, we first group the pairs that share the segment into groups of individuals that share the segment, declaring this as the IBD’s cohort. Then, for each *inferred* segment, we identify the ground truth *g* among all the ground truth IBDs in *G* that most closely matches the inferred segment, and we declare that this ground truth the match. We also have a threshold *t* and we allow *t/*2 of *g*’s base positions to be offset with *inferred*’s base positions. We refer the reader to Algorithm 2 for a simple pseudocode and Figure 3 for an example of an accepted segment).

**Figure 3.**
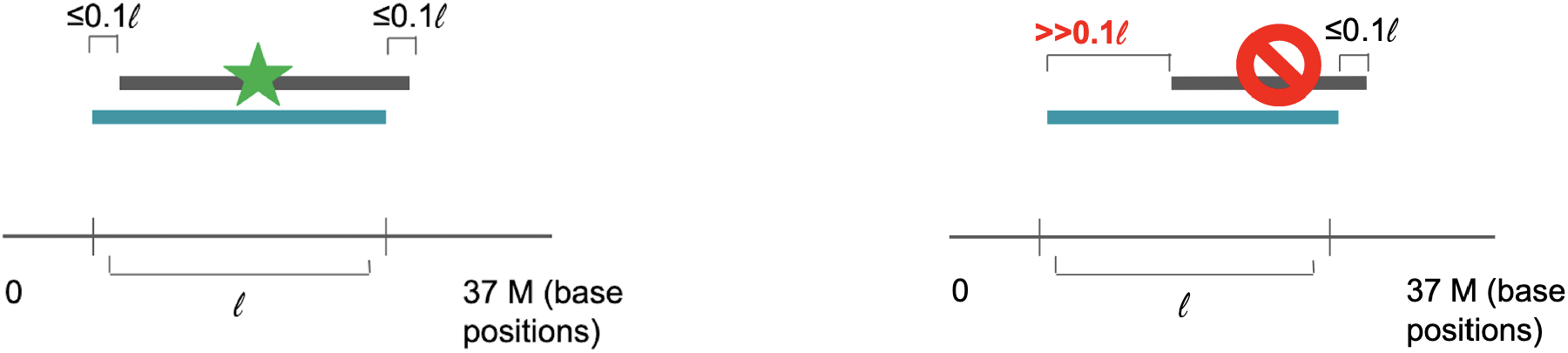
Matching inferred IBD segments (blue) with the ground truth (black). (left) Example of an instance where the inferred segment matches the ground truth. (right) Example where the inferred segment doesn’t match the ground truth. We refer to *t* as the IBD offset difference threshold.

#### Algorithm 2

Pseudocode for detecting segment overlap (between ground truth segments and inferred segments).

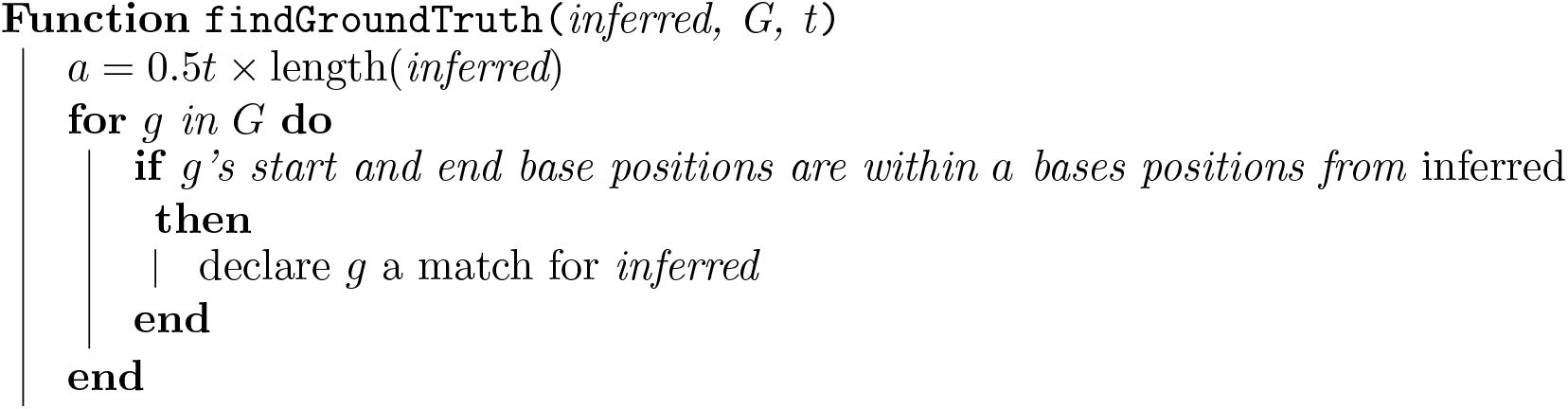

This approach can result in multiple ground truth matches for each inferred IBD. For now, we keep all of the ground truth matches in our analysis. We use this method to compare how IBD segments inferred by `phasedibd` correspond to individuals and to validate our kNN algorithm.

Next, we ran `phasedibd` on the simulated Amish pedigree with 1338 individuals with varying degrees of tolerance for errors in the IBD segments. `phasedibd` would return a set of IBD segment candidates which we use to compare against the true IBD segments that `Ped-sim` generated. We only select IBD segments inferred by `phasedibd` that have a significant amount of base position overlap with the true IBD segments generated by `Ped-sim`. We define the degree of overlap between two IBD segments by the fraction of matching base positions. We varied the amount of tolerance for offset difference from 0.001, 0.01, 0.015, 0.02, 0.03, 0.04 and 0.1. In other words, we accept inferred IBD segments that have at least 0.999, 0.99, 0.985, 0.98, 0.97, 0.96, and 0.9 fractional base positions overlap with any true IBD segment respectively. After filtering away the IBD segments that do not have sufficient overlap with the IBD segments from `Ped-sim`, we now possess cohorts of individuals corresponding to the IBD segments predicted by `phasedibd` that can be used as test cases against the cohorts corresponding to the real IBD segments generated by `Ped-sim`. We iterate through each of the valid IBD segments predicted by the two IBD detection software and run Algorithm 1 on the cohort that corresponds to the IBD segment to perform outlier removal for different parameters of *k* and *t*. Subsequently, we compare each of these processed cohorts to their original to determine the change in the number of individuals. We then determine how many of these individuals belonged to the corresponding true cohort from `Ped-sim`.

## Results

### Analysis of IBD detection software

We evaluated `phasedibd` and `GERMLINE`’s performance using the genotypes simulated by `Ped-sim` (using the Amish pedigree structure from [13]) as inputs for `GERMLINE` and `phasedibd`. We used the IBD segments simulated by `Ped-sim` as the ground truth to evaluate the correctness of the IBDs inferred by `GERMLINE` and `phasedibd` and observed some of the shortcomings of current IBD detection algorithms. The total length of IBD segments and the number of IBD segments inferred by `GERMLINE` both changed significantly when the genotyping error rate increased. On the other hand, the number of IBD segments inferred by `phasedibd` decreased slowly. When we increased the genotyping error rate, the total length of IBD segments inferred by `GERMLINE` decreased rapidly. `GERMLINE` inferred about 5 times fewer SNPs when the genotyping error rate was increased from 0.0 to 0.005, and almost 40 times fewer SNPs when the genotyping error rate went up to 0.01. When the genotyping error rate is higher, it seems that `GERMLINE` breaks IBD segments into smaller fragments due to their conservative hashing method.

On the other hand, `phasedibd` estimated only 20% fewer SNPs and about 7% fewer IBDs when the error rate went from 0.0 to 0.01. We believe that the `phasedibd`’s robust phase switch error correction explained this consistent performance. However, `phasedibd` overestimated the number of IBD segments and the total length of IBD segments when we compare its figures to what `Ped-sim` intended. `phasedibd` inferred twice as many total SNPs and three times as many IBDs between pairs as the ground truth. The same overestimation issue happened for `GERMLINE` in the ideal case when the genotyping error rate is set to 0.0. This indicates that both approaches were struggling with the problem of outputting too many false positive IBD segments.

Besides the disparities in the number and length of the IBD segments, we also observed a considerable number of false positives for the IBD segments inferred by current IBD detection algorithms. In Table 1, we list the IBD segments inferred between pairs of individuals and evaluate the number of True Positive (TP) SNPs and the number of TP IBD segments. In Table 1, when evaluating the IBD segments per SNP, we noticed that `phasedibd` correctly inferred many more base positions than `GERMLINE` at very similar Positive Predictive Values (PPV) i.e. 40-45%. `phasedibd` noticeably inferred many more TP IBDs when we look at the SNPs. However, for pairwise IBD count, `phasedibd` only outperformed `GERMLINE` when the genotyping error rate is 0.01. We believe that `phasedibd`, with its robust phase-switch error correction, is able to infer longer IBDs. Because these IBDs have more than 1% of their SNPs that do not match the ground truth IBDs, `phasedibd`’s TP segment count is lower than `GERMLINE`’s when the genotyping error rates are smaller. However, when the genotyping error rate is 0.01, despite much lower PPV (29.02% compared to 43.79%), `phasedibd` picks up 4 times as many true positive segments as `GERMLINE`. When we looked at the pairwise case segments, we noticed a concerning PPV for the pairwise case for `phasedibd` which fell far below that of `GERMLINE` for varying genotyping rates. We hypothesize that the FP base positions resulted in very different lengths for the inferred IBD segments when compared to ground truths. This was very likely the cause for a much lower PPV for IBD segment counts in some trials. However, when the genotyping error rate is at 0.01, `phasedibd` still outperformed `GERMLINE` in terms of TP count.

**Table 1.**
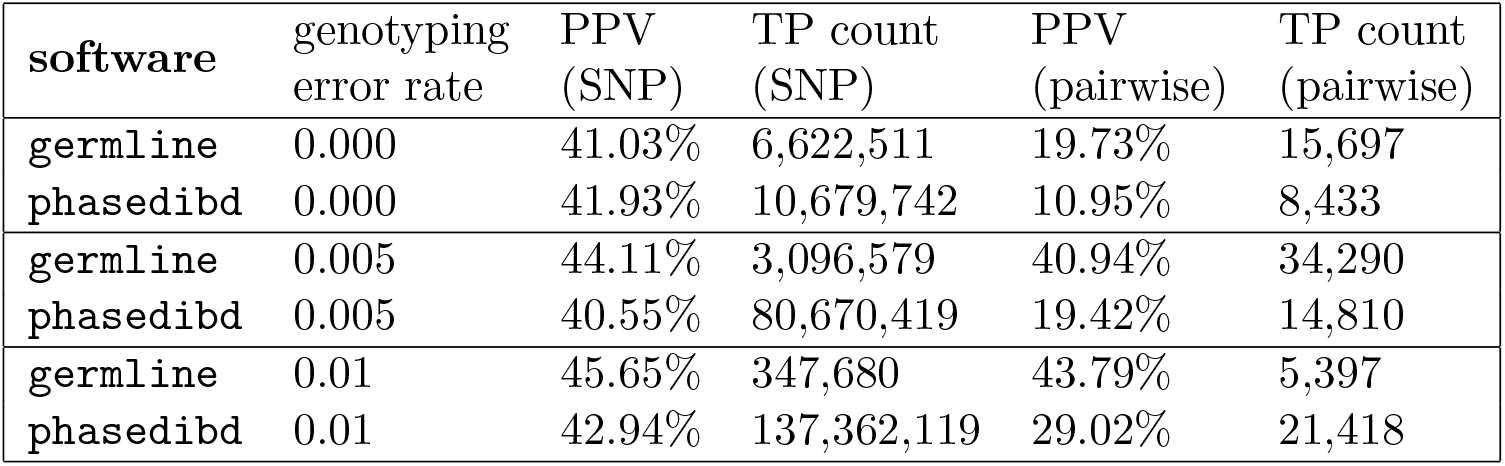
Comparison of `phasedibd` and `GERMLINE`’s performance for inferring IBDs of the genotyped descendants. We evaluated the performance on two metrics: Positive Predictive Value (PPV) and number of True Positives (TP). For per SNP evaluation, when an inferred IBD is observed between two individuals at certain SNPs and there exists a ground truth IBD at the same SNPs, we count this SNP as a TP, else we count this as a False Positive (FP). For pairwise evaluation, when an inferred IBD has a ground truth IBD that covers the same SNPs (we allow 1% of the SNPs in the ground truth IBD to mismatch with the inferred IBD), we increment TP Count, else, we increment FP count.

Overall, while `phasedibd` and `GERMLINE` both struggled with overestimating the amount of False Positive IBDs, `phasedibd` outperformed `GERMLINE` in trials with higher genotyping error rates. Therefore, we decided to improve upon this approaches performance by addressing the FP IBD segments.

As they were developed for general populations, we hypothesize that both `phasedibd` and `GERMLINE` overestimate the number of IBD segments in endogamous populations specifically. Hence, it is imperative to develop improved strategies to better distinguish IBD segments from IBS and background IBD in endogamous populations. As `phasedibd` performed better overal, we use its IBD segment calls as a starting point for our method.

### Inferring correct IBD cohorts

In our experiments, a cohort can have between 2 and up to 120 individuals that share an IBD segment. This highlights the applicability of our approach for both common and rare IBD segments. Correctly identifying these cohorts can bring us closer to having accurate IBD calls for downstream analysis. In Figure 4, we can see that when the genotyping error rate was increased, `phasedibd` built cohorts with more FP individuals; in other words, there was a higher number of IBD segments with at least 50% of their cohorts being FP individuals. We also observed that `phasedibd` performs worse when inferring IBD cohorts at higher genotyping error rates. In Figure 4, we provided histogram data visualization for the distribution of false positive individuals in the different cohorts under varying genotyping error rates. Figure 4 showed that `phasedibd` had a consistently larger number of cohorts with different percentages of false positive individuals in those cohorts. As the genotyping error rate increased, we noticed an increase in the number of cohorts with varying percentages of false positive individuals. For example, when the genotyping error rate was increased from 0.0 to 0.005, we noticed that the number of cohorts for the different false positive percentages doubled on average. However, when the genotyping error rate was further increased to 0.01, the number of cohorts decreased and we started to observe an increase in the number of cohorts with larger percentages of their cohort consisting of FP individuals.

**Figure 4.**
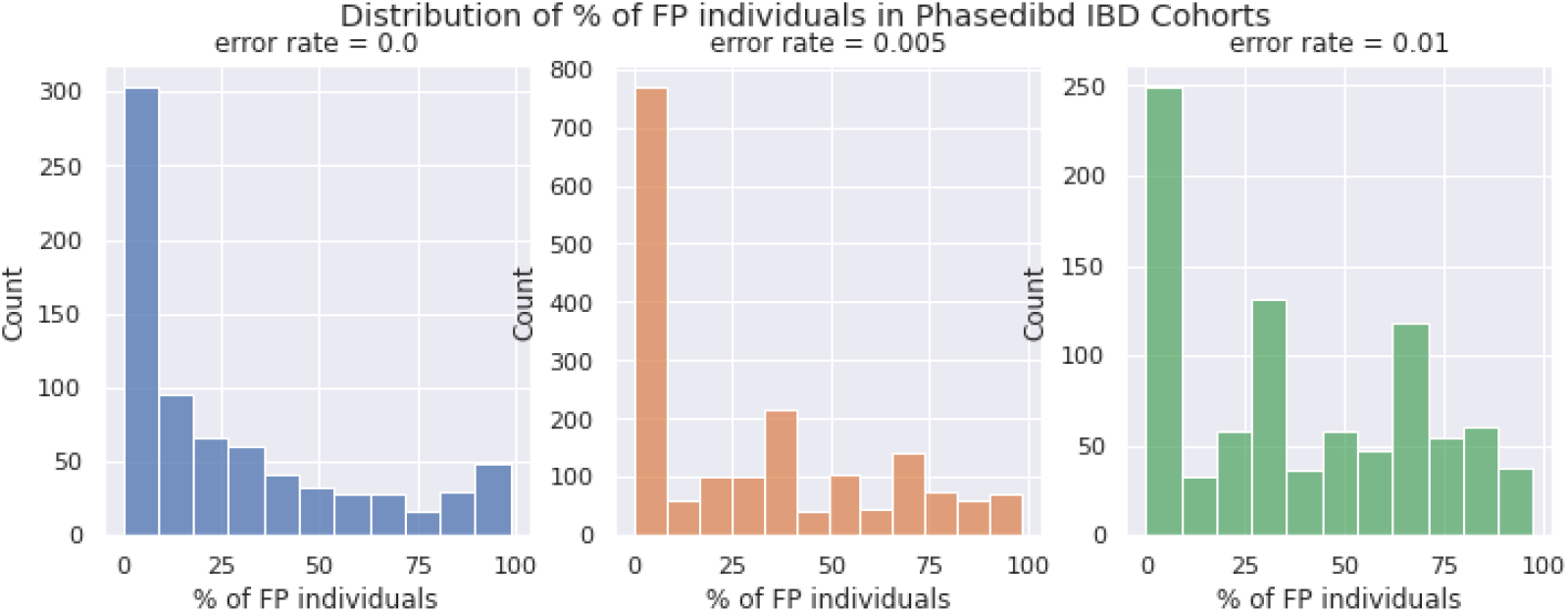
Distribution of the percentage of false positive individuals in `phasedibd` IBD cohorts for different genotyping error rates.

### Results of the kNN algorithm

As mentioned in the Validation section, we ran the kNN algorithm on the cohorts that corresponded to the IBD segments inferred by `phasedibd`. Because we varied the genotyping error rate over 0.0, 0.005, 0.01 and the IBD segment offset difference over 0.001, 0.01, 0.015, 0.02, 0.03, 0.04 we have 36 different configurations. For each of the configurations, we tried the following different *t*_soft_ values: 0.00625, 0.0125, 0.025, 0.05, 0.1, 0.2, 0.4, 0.8, 1.0. The *t*_soft_ values are chosen to provide the maximum variation in the *T*_final_ used for determining the outliers. We also varied the *k* value over 2, 8, 14, 20, 26, 32, 38, 44, and 50. We let *t*_hard_ be 0.0 for all subsequent results unless explicitly stated.

We compared the PPV values before and after running the kNN algorithm on the cohort. The original PPV values are computed by dividing the number of true positive members in the cohort over the sum of true positive and false positive members. For each of the three different genotyping error rates, we calculated the original PPV by averaging the original PPV across the different IBD segments inferred from the genetic data. To compute the PPV after running the kNN algorithm, we first computed the PPV across each of the kNN algorithm parameters i.e. k and *t*_soft_ for every IBD segment in our test cases. In the final results we compute the average PPV values across these test cases (for fixed *k* and *t*_soft_).

We now discuss how our kNN algorithm performs as we vary the IBD offset difference. In Figure 5, we see how the PPV values for both before and after running the kNN algorithm decrease as the IBD offset difference rate increases from 0.0 to 0.01. This is expected because more FP individuals would be able to infiltrate the cohorts if we loosen the requirements for IBD segments. However, we also note that kNN consistently outputs better PPV values than the original PPV. Moreover, the difference between the original PPV, the PPV value that corresponds to the cohorts before we run the kNN algorithm on them, and the kNN PPV, the PPV value that corresponds to the cohorts after we run the kNN algorithm on them, widens as the IBD offset difference rate increases. In other words, we expect the kNN algorithm to provide a much more significant improvement in the accuracy of cohort identification when we operate in the context where IBD offset difference is high.

**Figure 5.**
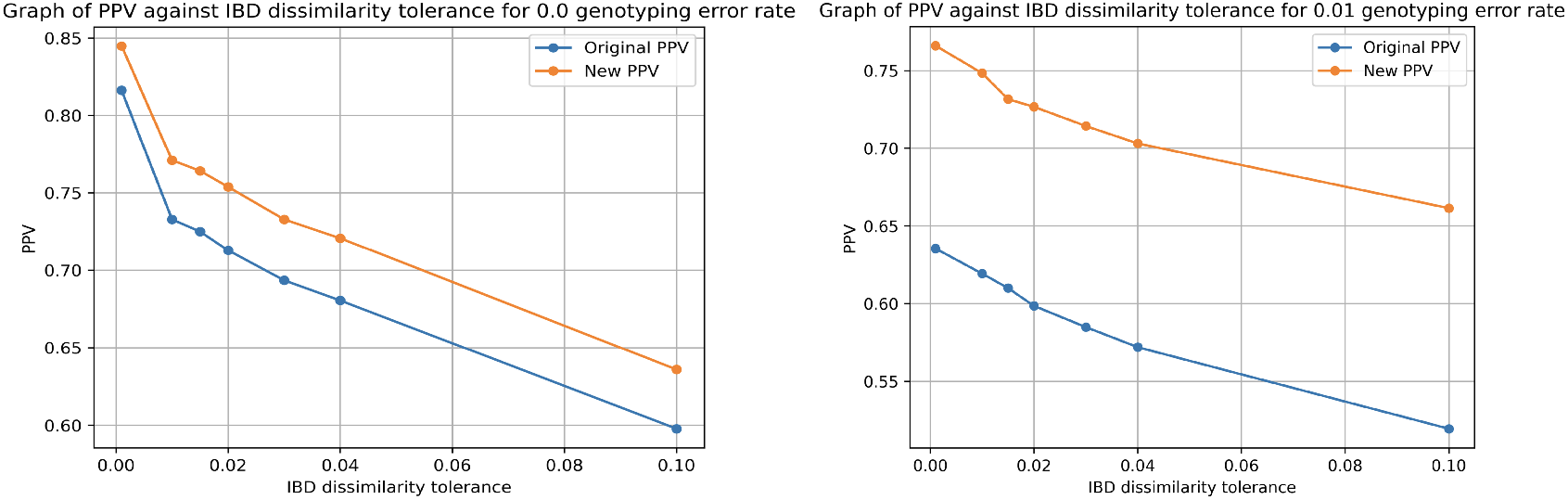
PPV against IBD offset difference for 0.0 genotyping error rate (left) vs. 0.01 genotyping error rate (right)

### Varying the fraction of cohort members to keep

While the above results appear to be very promising, we observed that the optimal kNN algorithm parameters do not necessarily give us realistic cohorts at the end of the false positive removal steps. We observe that a large proportion of the cohorts only contained two individuals after running the kNN algorithm on them. While this contributed to a marked improvement in the number of true positive individuals in the post-processed cohort, cohorts that contain only two individuals would not be optimally useful for ancestral haplotype reconstruction or studies of heritable diseases. This is related to the fact that the PPV values can be easily boosted when the cohorts are extremely small. Thus, we vary the *t*_hard_ values to ensure that we have a meaningful number of individuals in a cohort to better interpret the performance of our algorithm alongside the PPV metric.

We detail the performance of Algorithm 1 as we vary the parameter *t*_hard_ over the values of 0.0, 0.1, 0.2, 0.3, 0.4, and 0.5. In other words, we tracked the results of Algorithm 1 as we impose the condition of keeping at least 0%, 10%, 20%, 30%, 40%, and 50% of the cohort after running the algorithm. Note that our results in the previous subsection correspond to the case where *t*_hard_ is equal to 0.0.

From Figures 6a and 6b, we note that as the *t*_hard_ values were increased from 0.0 to 0.5, the improvement in PPV brought about by the kNN algorithm decreased. However, we note that the decrease in the improvement in PPV was not substantial except for the case where we have 0.0 genotyping error rate and 0.001 IBD offset difference. Even when we enforce the requirement to keep at least 50% of the cohort, we still observe a non-trivial improvement in PPV i.e. the ratio of true positives in the cohort still improved after running the kNN algorithm on the cohorts. In fact, we observe that constraining ourselves to keep a significant minimum fraction of the cohort barely impacts the PPV gains we obtain from running the kNN algorithm most of the time. Thus, we conclude that adding in the additional parameter *t*_hard_ would resolve the issue of cohorts only containing 2 individuals and still achieve an improvement in the PPV that is almost as good as the case where we do not have the *t*_hard_ parameter.

**Figure 6.**
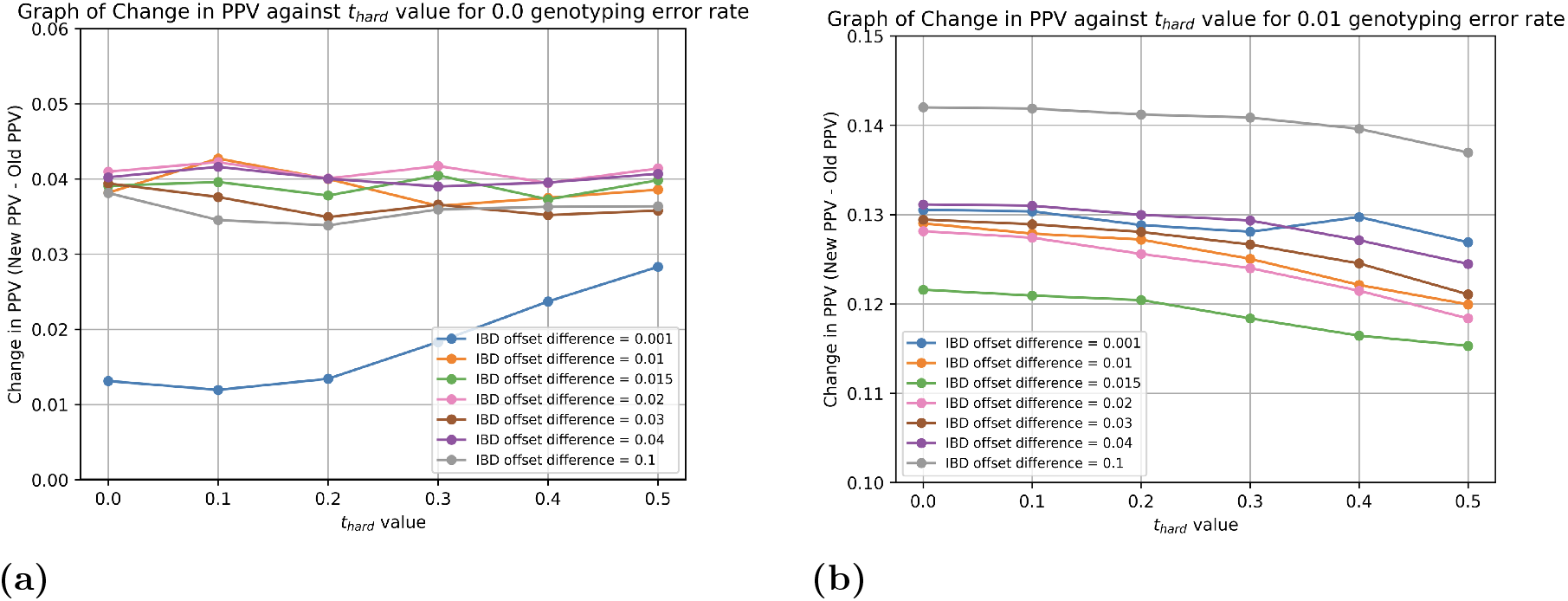
Average Change in PPV for `phasedibd` with different genotyping error rates against *t*_hard_ values

## Discussion

In this work, we propose a new method to refine IBD segment calls in endogamous populations, where false postitive due to IBS and background IBD are common. We provide a formal algorithmic approach to the problem of removing outliers in the inferred cohort of individuals that correspond to an IBD segment detected by `phasedibd`. We evaluated the performance of our algorithm on an endogamous Amish pedigree with genomic data simulated by `Ped-sim` with varying genotyping error rates. Genetic studies of Old Order Amish are ongoing, with several groups studying common and rare diseases and traits [19, 23, 24]. Using our kNN algorithm, it is possible to enhance the performance of IBD detection software by improving the accuracy of cohort determination. In other words, we are better able to determine the group of individuals who have inherited a particular IBD segment. This should help clinicians and researchers better determine which haplotypes contribute to different genetic disorders in endogamous pedigrees. We believe our algorithm also has the potential to improve ancestral haplotype reconstruction by helping improve the assignment of IBD segments to ancestors. While the results we have obtained are promising, they are not sufficiently general for us to make the claim that they work for all endogamous populations, given our analysis was restricted to an Amish pedigree.

Some readers might wonder why the shortest path between individuals in a pedigree tree is chosen out of all possible paths to be the distance metric. The concern is valid since the purpose of Algorithm 1 is to provide us with a means to develop a more conservative estimation of the cohort by removing outliers. It would make sense to perhaps consider paths that are longer. However, we note that the longest path between two individuals would be the trivial path from one individual to a founder of the pedigree and then to the other individual. Hence, the longest paths do not necessarily capture information regarding whether individuals belong to a cohort or not. In fact, the shortest path should carry the most weight since the existence of a short path between two individuals immediately suggests a high probability of sharing genetic segments. However, we do acknowledge that analyzing multiple non-trivial paths between individuals could provide a much more reliable set of statistics that could be used to develop a better distance metric. An example would be the kinship coefficient score that can be computed by KING [25]. Kinship coefficients consider the pedigree structure in a much more comprehensive way. Thus, it could potentially be a better distance metric for us to use for the kNN algorithm.

Another possible extension to our work is the estimation of the optimal kNN parameters for different cohort contexts. As mentioned in the results section, the kNN parameters required for optimal false positive removal where we maintain minimal false negatives tend to vary for different IBD segments and cohort sizes. Thus, it could be useful to train a machine learning model to estimate the optimal set of parameters for the kNN algorithm to guarantee a high efficacy rate for false positive removal with our kNN algorithm. To do that effectively, it might be necessary to simulate more different endogamous pedigrees using `Ped-sim`.

There are also many other algorithmic modifications that we can explore for our kNN algorithm. As of now, we are only using the shortest path between two individuals as a distance metric to facilitate the outlier detection process in the kNN algorithm. However, it is possible for us to derive information about the cohorts that are far richer than the average distance between an individual and its *k* nearest neighbors. One possible insight that is specific to endogamous populations is the presence of loops in the pedigree tree. It is possible that detecting loops within a cohort of individuals might help improve the accuracy of outlier detection. Individuals related by a loop within the cohort are much more likely to share an IBD segment. Hence, detecting loops within the cohort may be a productive pre-processing step for the kNN algorithm.

Another statistic worth analyzing is how the kNN density varies across the members within the cohort. Right now, the design of the kNN algorithm requires a jump in the kNN density and uses that jump to determine if an individual in the cohort is an outlier. Future work could analyze how the kNN density varies from true positives to false positives to better understand how we can determine the threshold for determining false positives.

